# Moments of the length of sample genealogy and their applications ☆

**DOI:** 10.1101/2025.11.21.689774

**Authors:** Yun-Xin Fu

## Abstract

Underlying many important statistical properties of a sample of DNA sequences is the total branch length *L* of the sample’s genealogy. A classic example is the number *K* of mutations on the genealogy, whose expectation and variance are simple functions of the expectation and variance of *L*. However, higher moments of *L* and related quantities have received relatively little attention despite their potential utility. This paper systematically investigates the properties of *L*, its relationship with the total branch length of a certain number of descendants, and demonstrates their usefulness through several applications under the constant-in-state model, which is an extension of the Wright–Fisher model with constant effective population size. Specifically, the recurrence relations for the power moments of *L* as well as for the expectations of products between powers of *L* and branch length of specified sizes are derived. A closed-form expression for the power moments of *L* is also obtained. These results allow examination of the large-sample behavior of *L* through its skewness and kurtosis, revealing that *L* does not satisfy asymptotic normality under the constant-size Wright–Fisher model, although approximate normality emerges in rapidly growing populations. Moreover, the power moments of *L* provide a more straightforward route to deriving higher moments of *K* and yield a novel approach for computing the distribution of *K*. The associated mixed moments similarly lead to a novel method for calculating the probability of having a single mutation of a specified number of descendants.

---

Coalescent times are fundamental components of the coalescent process (Kingman, 1982b,a), as they, together with the mutation rate, largely determine both the amount and the pattern of genetic variation observed in a sample of DNA sequences. However, most summary statistics used to characterize sample polymorphism do not relate directly or simply to coalescent times. Instead, they typically depend on intermediate summary statistics of these times, one of the most important being the total branch length *L* of the sample’s genealogy. A classic example is the number *K* of mutations on the genealogy, which equals the number of polymorphic (segregating) sites under the infinite-sites model (Ewens, 2004). For a sample of sequences from a single locus without recombination, Watterson (1975) derived the expectation and variance of *K* as

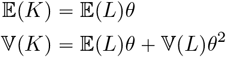

where *θ* = 4*Nμ* with *N* being the effective size of the population and *μ* being the mutation rate per sequence per generation.

To date, the use of *L* has been driven mainly by its expectation, variance, and covariances with other summary statistics. This is partly because many applications require no more than second moments, but it also reflects the limited understanding of *L* itself, including the absence of efficient methods for handling its higher moments. As statistical inference based on sample polymorphism becomes increasingly sophisticated, the need to compute the probability of specific polymorphism patterns has grown—and such probabilities are closely tied to the higher moments of *L*.

The purpose of this paper is to systematically investigate higher moments related to *L* and to illustrate their applications for a sample of DNA sequences from a single locus in a population evolving according to the constant-in-state model (Fu, 2025a,b), which includes the Wright– Fisher model with constant effective population size as a special case. Specifically, we derive both recurrence relations and a closed-form formula for the power moments of *L*, which in turn facilitate the derivation of recurrence relations for the expectations of products between the length of branches with a given number of descendants and powers of *L*. These results yield concise derivations of the probability and moments of the number of mutations in the sample genealogy, the probability that a single muta-tion has a specified number of descendants, and a detailed examination of the skewness and kurtosis of *L*. The latter analysis shows that *L* departs substantially from normality under constant population size, whereas it agrees well with normality under rapid population expansion.

## 1. Constant-in-state model and summary statistics of coalescent times

Consider a sample of *n* sequences of a non-recombining locus from a single population evolving according to the Wright–Fisher model (Ewens, 2004). Under the Kingman coalescent (Kingman, 1982b,a) with a constant effective population size *N*, the *i*-th coalescent time, which is the waiting time for *i* ancestral lineages to coalesce into *i* −

1 lineages, follows an exponential distribution with rate parameter *i* (*i* − 1), where one unit of time corresponds to 4*N* generations. Although the classical derivation of the Kingman coalescent assumes *n* ≪ *N* under the Wright– Fisher model, it is known to be robust under alternative models (such as the Moran model) that do not impose this restriction. Therefore, we assume that *t*_*i*_ follows an exponential distribution with rate for all *i* ⩽ *N* . Under the Kingman coalescent, the *t*_*i*_ are mutually independent.

The constant-in-state model (Fu, 2025a,b) maintains the independence of coalescent times while allowing the effective size *N* to vary among the states of the coalescent process. Due to its flexibility for modeling non-constant population dynamics and its mathematical tractability, it has found successful applications in the inference of population history (Pybus et al., 2000; Liu and Fu, 2015, 2020).

For a sample of size *n*, let *N*_*i*_ be the effective population size in state *i* (*i* = 2, …, *n*), and let *v*_*i* − 1_ *N*_*i*_ /*N*, where *N* is the effective population size at a reference state (without loss of generality, state 2 is chosen as the reference). Under the constant-in-state model, the *i*-coalescent time *t*_*i*_ therefore follows an exponential distribution with parameter *i* (*i* = 1 /*v*_*i* − 1_, where time is measured in units of 4*N* generations. A constant-in-state model is fully specified by the row vector

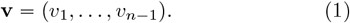

For brevity of notation, we adopt the following conventions: 1) if *v*_*i* + 1_ is not specified explicitly, then it is assumed that *v*_*i* + 1_ = *v*_*i*_; 2) if *v*_*i*_ = *f* (*i*) for some function *f* ( *i*), then **v** will be written as (*f* (*i*) ) ; and 3) *a* (*k*) represents *k* consecutive *v*_*i*_ *a*. Following these conventions, a population with constant effective size is specified as **v** = (1) .

The constant-in-state model is capable of mimicking a wide range of population dynamics, including population bottlenecks and growth models. For example, **v** = (1(5), 0.05(2), 1) represents a bottleneck scenario, while for *α* > 0, **v** = (*i*^*α*^) represents population growth. The follow-ing three examples illustrate growth at different speeds:

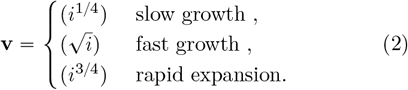

It follows that *N*_100_ under these three models is 3.1, 10, and 31.6 times, respectively, larger than the population at state 2, justifying their descriptive labels.

The coalescent process yields a genealogy of the sample, from which various summary statistics of the coalescent times can be defined and computed. Examples include the age of the most recent common ancestor (MRCA) of the sample, the length of the two immediate descendant branches of the MRCA, the mean time separating two sequences, the total branch length of the genealogy, the total length of branches with exactly *i* descendants in the sample, and the lengths of external and internal branches. The expectations, variances, and covariances of these and other summary statistics of coalescent times have been explored in the literature (Watterson, 1975; Tajima, 1983, 1989; Fu and Li, 1993; Fu, 1995, 2022; Uyenoyama, 1997; Alimpiev and Rosenberg, 2022).

The total branch length *L* of the genealogy is the sum of the lengths of all branches in the genealogy. By segmenting the genealogy according to coalescent states, each branch consists of one or more segments, which are referred to as lineage segments(or simply lineages). For example, the external branch leading to seq. 4 in Fig. 1 consists of two lineages labeled 5 and 9, while the longer branch descending from the MRCA consists of lineages 1 and 3. It is easy to see that *L* can be expressed as

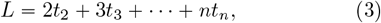

which is independent of the genealogical relationships among the sequences (Fig. 1).

**Fig. 1.**
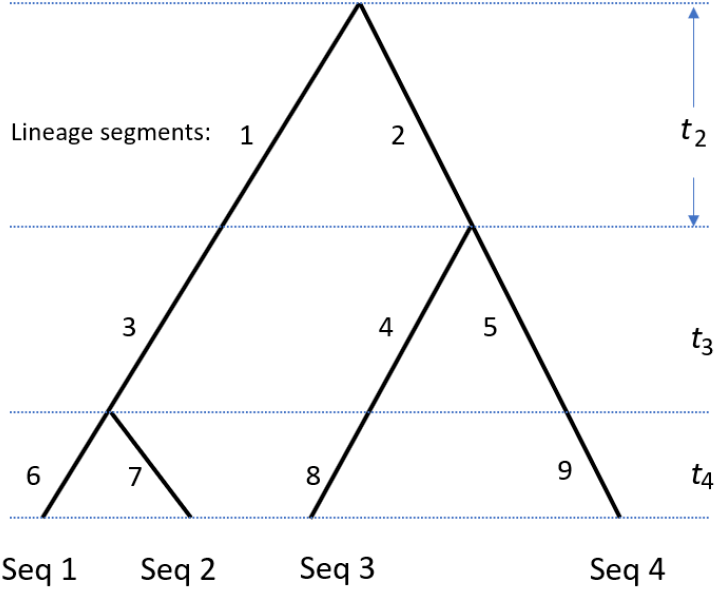
Illustration of *L* and *L*_*i*_ in a genealogy of four sequences. Lineages are numbered from 1 to 9 leading to 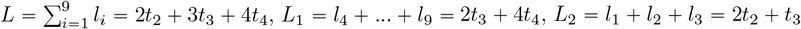 and *L*_*3*_.

Each branch (and its constituent lineages) in a genealogy can be classified by the number of descendants it has in the sample. A branch that has exactly *i* descendant sequences is said to be of size *i*, as are the mutations occurring on such a branch (Fu 1995). Therefore, each branch (and its constituent lineages) has a size between 1 and *n* − 1. The total branch length of size *i* is denoted by *L*_*i*_, which can be determined for a given sample genealogy (Fig. 1), but can also be expressed in general as follows. Suppose the *k* lineages of state *k* are labeled (for example, from 1 to *k*), and define, for the *l*-th lineage, an indicator variable *χ*_*kl*_ (*i*) that takes the value 1 if the lineage is of size *i*, and 0 otherwise. Then

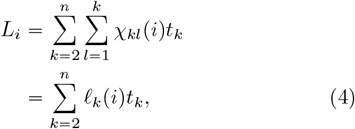

where 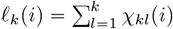 is the number of lineages of size *i* in state *k*. Also note that the *L*_*i*_ are mutually exclusive and satisfy the relationship

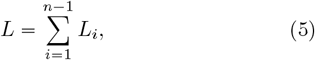

Throughout the paper, 𝔼 denotes the expectation. Since 𝔼 (*L*^*m*^) and 𝔼 (*L*_*i*_*L*^*m*^) are our primary focus, we define

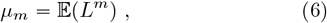

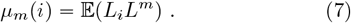

For a population of constant size, it is well known (Watterson, 1975) that

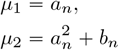

where

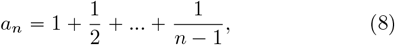

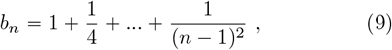

while from Fu (1995), we have that

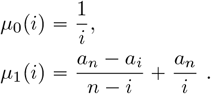

For a constant-in-state model **v**, define a generalized zeta function for *i* > 0

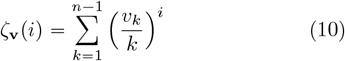

Note that the *ζ* (*i*) defined in Fu (2025a) for the derivation of moments of *K* is equal to *ζ*_**v**_ (*i*) *θ*^*i*^. The current notation, although less concise, is preferred because it relates to, but is distinct from, the *ζ* (*s*) commonly used for the Euler– Riemann zeta function (Abramowitz and Stegun, 1972). Since *ζ*_**v**_(1) appears frequently in the paper, it may be shortened to *ζ*_**v**_ for brevity. It follows that *a*_*n*_ = *ζ*_(1)_(1) and *b*_*n*_ = *ζ*_(1)_(2). Throughout the paper, the *k*-th falling factorial of *x* is denoted by (*x*)_*k*_, that is,

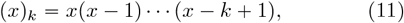

with the convention that (*x*)_0_ = 1.

## 2. Moments of *L*

Let 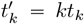, which is the total time of the sample genealogy at state *k*. Then

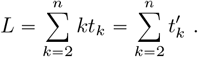

It follows that 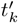 is exponentially distributed with parameter *λ*_*k*_ = (*k* − 1){*v*_*k*−1_. Therefore

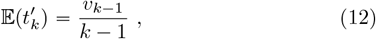

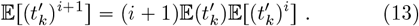

Since 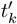 are mutually independent, the moment generating function (mgf) of *L* is

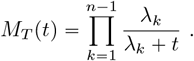

In principle, the standard method, which differentiates the moment generating function (mgf) and then sets *t* = 0, can be used to obtain *μ*_*m*_. This approach allows the derivation of the first few moments, but generalizing to large values of *m* is difficult, similar to the derivation of higher moments of *K* (Fu, 2025a). A more efficient alternative is to utilize the cumulants of *L*, whose generating function is ln *M*_*T*_ ( *t*) . Recurrent relationships exist for deriving power moments from cumulants (e.g., Smith (1995)). Since we aim to examine not only *μ*_*m*_ but also *μ*_*m*_ (*i*), which is more complex, we opt for a more direct approach and first prove the following intermediate result.

### Lemma.

*For a random sample of n sequences from a population evolving according to a constant-in-state model* ***v***. *Then for m* ⩾ 0 *and* 2 ⩽ *k* < *n*

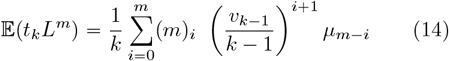

*where* (*m*)_*i*_ *is the i-th falling factorial of m*.

### Proof.

Let 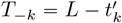. Then

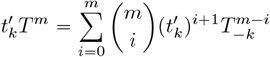

Since 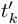 and *T* _− *k*_ are independent, it follows from Eq.(12) and Eq.(13) that

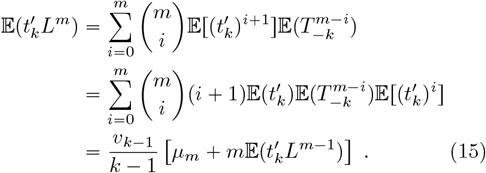

By repeatedly substituting Eq.(15) for 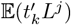, we arrive at a recurrence equation for 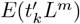 which is e)uivalent to (14).|

The lemma allows us to obtain a recurrence e)uation for *μ*_*m*_ as follows

### Theorem 1.

*For a random sample of n sequences from a population evolving according to a constant-in-state model* ***v***. *Then for m* > 0

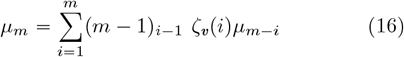

*where ζ*_***v***_(*i*) *is defined by Eq*.(*10*) *and* (*m* − 1)_*i*−1_ *is the* (*i* − 1)*th falling factorial of m* − 1.

### Proof.

Since 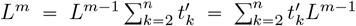, it follows from the Lemma that for *m* > 0

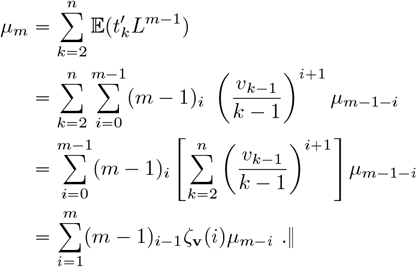

The theorem provides an efficient way to compute *μ*_*m*_ sequentially, but it is useful to have a close-form expression for *μ*_*m*_, particularly for theoretical study. The following corollary provides such a solution by expressing *μ*_*m*_ as the sum of products related to *ζ*_**v**_, which requires partitioning *m* as a sum of smaller integers. A partition of *m* can be uniquely represented by multiplicity notation

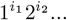

where *i*_*j*_ is the number of “j” appeared in the parts, which satisfies

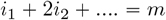

Notation 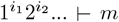 means that 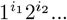 is a partition of *m*. Order of parts in a partition does not matter but it is generally arranged in descending order.

### Corollary 1.

*For a random sample of n sequences from a population evolving according to a constant-in-state model* ***v***. *Then for m* > 0

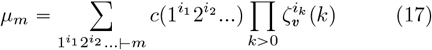

*where the summation is taken over all integer partitions of m, ζ*_***v***_(*k*) *is defined by* (*10*) *and*

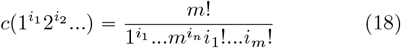

### Proof.

We will use induction to prove the theorem. Since *μ*_1_ = *ζ*_**v**_(1), and there is only one partition of 1 with *c*(1^1^) = 1, equation (17) is thus true. Suppose it is true for up to *m* − 1. Then due to Theorem 1

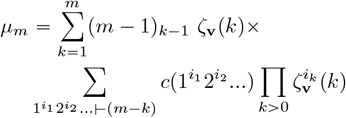

Since adding a ^′^ *k*^′^ to partition 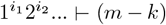 leads to a partition 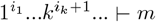 with the relationship

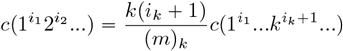

and every partition of *m* will lead to partitions of *m* − *k* when one of its part ^′^ *k*^′^ is removed. Grouping terms according to the partitions of *m* leads to

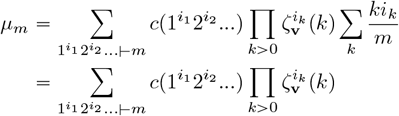

Therefore Eq.(17) is also trure for *m*. ‖

It should be noted that 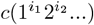 is known as the number of ways to arrange *m* different objects into cycle type defined by partition 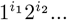 (Abramowitz and Stegun, 1972; Riordan, 1958). This number was found in Ewens’ sampling formula (Ewens, 1972; Karlin and McGregor, 1972) and recently found to be in the distribution of the number *K* of mutations in the sample genealogy (Fu, 2025b).

The numbers of integer partitions for *m* = 1, …6 are, respectively, 1, 2, 3, 5, 7 and 11 (e.g., Table 1 of Fu (2025b)), which indicate the number of different terms in *μ*_*m*_ for *m* ⩽ 6. Take *μ*_3_ for example. The three integer partitions are 3^1^, 1^1^2^1^ and 1^3^ respectively, while

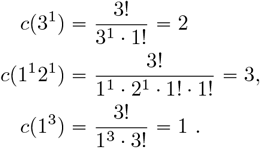

**Table 1.** The minimum *h* such that ℙ_*n*_ (*K, h*) has less than 1% relative error for a sample of size 100 from a population of constant size.

| K | h | $\mathbb{P}_n(K)$ | K | h | $\mathbb{P}_n(K)$ |
| --- | --- | --- | --- | --- | --- |
| $\theta = 0.1, \mathbf{v} = (1)$ | | | $\theta = 0.5, \mathbf{v} = (1)$ | | |
| 1 | 3 | $3.01 \times 10^{-1}$ | 1 | 18 | $2.03 \times 10^{-1}$ |
| 2 | 4 | $8.01 \times 10^{-2}$ | 2 | 20 | $2.41 \times 10^{-1}$ |
| 5 | 4 | $2.90 \times 10^{-4}$ | 5 | 29 | $7.27 \times 10^{-2}$ |
| 10 | 6 | $3.28 \times 10^{-9}$ | 10 | 48 | $9.03 \times 10^{-4}$ |
| 20 | 10 | $1.34 \times 10^{-19}$ | 20 | 90 | $1.89 \times 10^{-8}$ |
| $\theta = 0.1, \mathbf{v} = (i^{1/4})$ | | | $\theta = 0.5, \mathbf{v} = (i^{1/4})$ | | |
| 1 | 5 | $3.62 \times 10^{-1}$ | 1 | 27 | $5.36 \times 10^{-2}$ |
| 2 | 5 | $1.66 \times 10^{-1}$ | 2 | 29 | $1.13 \times 10^{-1}$ |
| 5 | 5 | $2.56 \times 10^{-3}$ | 5 | 36 | $1.62 \times 10^{-1}$ |
| 10 | 6 | $1.40 \times 10^{-7}$ | 10 | 51 | $1.46 \times 10^{-2}$ |
| 20 | 10 | $1.23 \times 10^{-17}$ | 20 | 90 | $1.49 \times 10^{-6}$ |

$\mathbb{P}_n(K)$ is computed by Eq.(11) of Fu (2025b).

Therefore according to Corollary 1

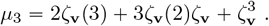

It is straightforward to derive the concise expressions of *μ*_*m*_ for relative small *m*, and the collection for *m* ⩽ 6 is as follows

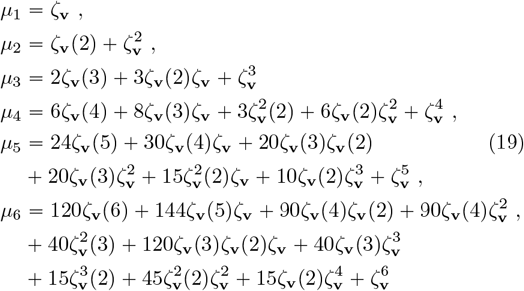

which allows one to study the commonly used central moments. Since

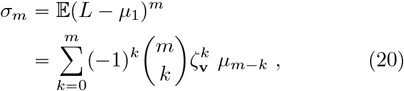

it is straightforward to show that

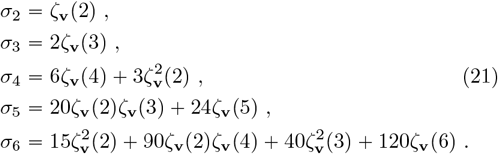

Fig. 2 plots the logarithm of *μ*_*m*_ under models **v** = (1) and 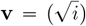. In the log scale, moments differ little for different sam(le sizes under **v** = (1). However, the difference for the model 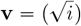 is considerable. In each case, linearity is observed for large *m*, indicating *μ*_*m*_ increase exponentially for large *m*.

**Fig. 2.**
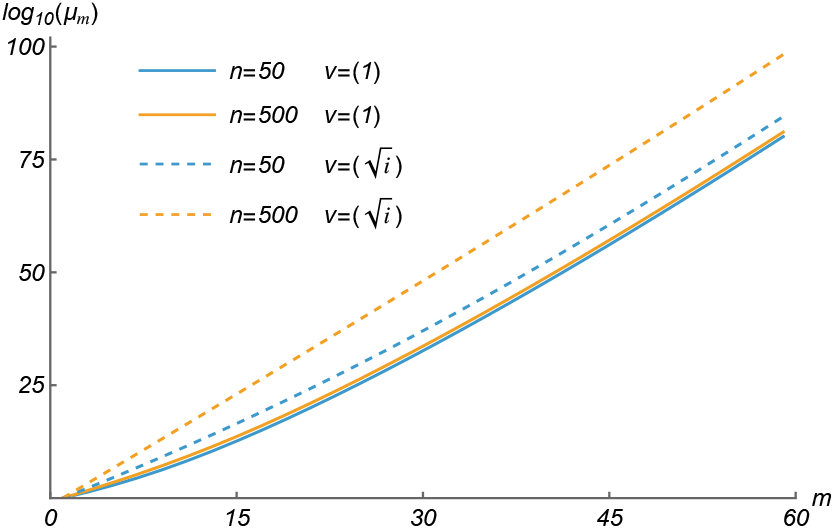
*log*_10_(*μ*_*m*_) under the constant population size model (**v** = p1)) and fast growing model 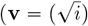 for two sample size *n* = 50 and 500.

## 3. Expectation of *L*_*i*_*L*^*m*^

The expectation of *L*_*i*_*L*^*m*^ is useful for finding the probability of a single mutation of size *i*, which will be explored in a later section. From (4) it follows (Fu, 1995) that

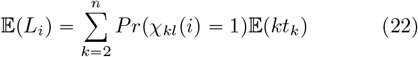

where *Pr* (*χ*_*kl*_ *i* = 1) is the probability, *p*_*n*_ (*k, i* ),that a randomly selected lineage from state *k* is of size *i*. Fu (1995) shows that

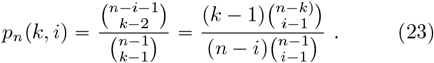

Define another zeta function for 0 < *i* < *n* and *j* ⩾ 0

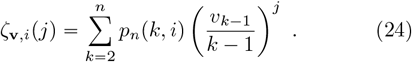

Then *ζ*_**v**,*i*_ (0) is the expected number of lineages of size *i* when one lineage is randomly selected from each state, while *ζ*_**v**,*i*_(1) = *E*(*L*_*i*_). Furthermore

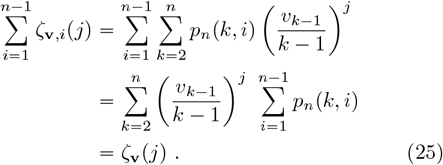

Therefore, *ζ*_**v**,*i*_ (*j*) ‘s partition *ζ*_**v**_ (*j*) into *n* − 1 positive components.

From Eq.(4), we can write

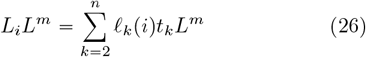

where ℓ_*k*_ (*i*) is the number of lineages at state *k* that is of size *i*. Therefore

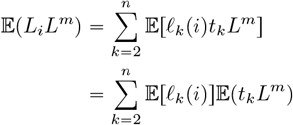

where the second equality is due to that each coalescent is between two random lineages and thus is independent of the coalescent times. Since 𝔼[ℓ_*k*_(*i*)] = *kp*_*n*_(*k, i*) while 𝔼(*t*_*k*_*L*^*m*^) is given by Eq.(16), it follows that

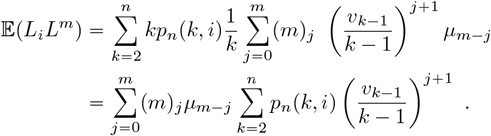

We therefore have the following theorem

### Theorem 2.

*For a random sample of n sequences from a population evolving according to a constant-in-state model* ***v***. *Then for m* ⩾ 0

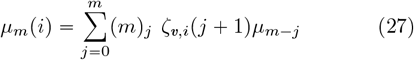

*where ζ*_***v***,*i*_ (*j*) *is defined by Eq*.(*23*) *and m* _*j*_ *is the jth falling factorial of m*.

Together with (19) the theorem immediately leads to

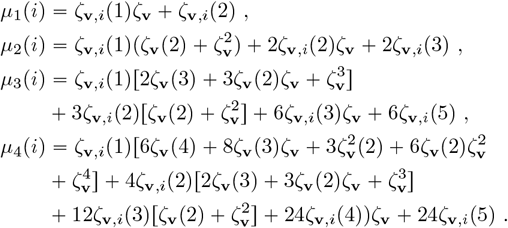

Also due to Eq.(5), we have for *m* > 0

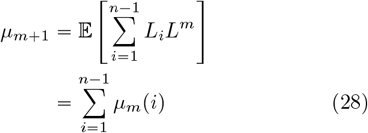

which can also be verified by summing up Eq.(27), resulting in the right-hand side of the equality equal to *μ*_*m*+1_ due to Eq.(25) and Theorem 1.

Since it is necessary to calculate *ζ*_**v**,*i*_(*j*) for evaluating *μ*_*m*_(*i*), it is useful to explore efficient way for its computation. A simplified expression of *ζ*_**v**,*i*_(*j*) can be derived under model **v** = (1). Because of Eq.(23), it follows that

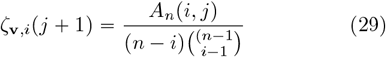

where

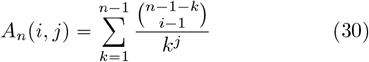

It follows that *A*_*n*_ (1, *j*) = *ζ*_**v**_ (*j*) . Because 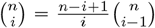 we have the following recurrence relationship

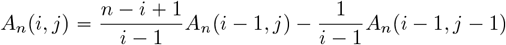

Continuing to replace the first term by this recurrence equation leads to

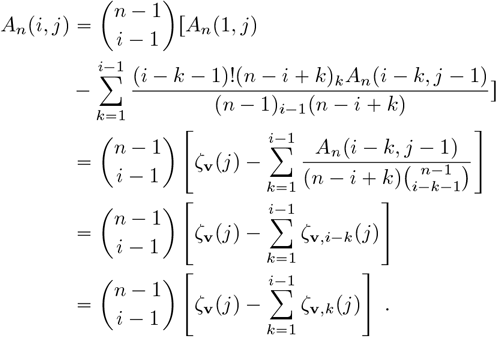

Therefore due to (25) we have

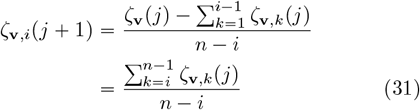

with the initial condition *ζ*_**v**,*i*_(1) = 1/*i*. Therefore *ζ*_**v**,*i*_(*j* + 1) can be obtained as the average of *ζ*_**v**,*k*_(*j*) for *k* ⩾ *i*.

Fig. 3 plots log_10_ [*μ*_*m*_ (*i*)] for several values of *i* under both the constancy and fast growth model. The overall trend for each case is similar to that for log_10_ (*μm*) . However, the difference between different *i* is marginal, suggesting that *L*^*m*^ dominates the value of *μ*_*m*_(*i*).

**Fig. 3.**
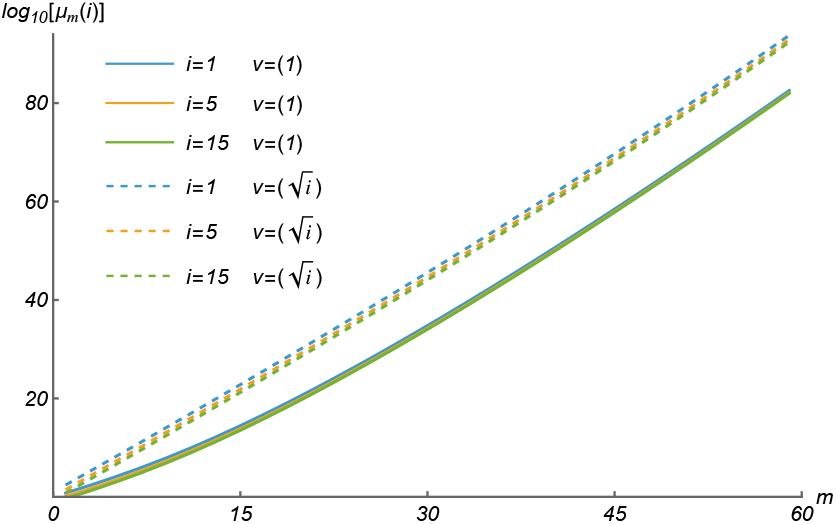
Examples of log [*μ*_*m*_(*i*)] under constant population size (**v** = (1)) and expanding model 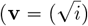 for *n* = 200.

## 4. Skewness and kurtosis of *L*

The higher moments of *L* allows one to examine two important aspects of *L*, its symmetry and tailedness. The former is measured by the skewness *γ*_1_ and the latter by (excess) kurtosis *γ*_2_, which are respectively

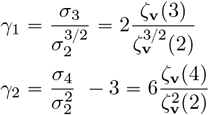

Under the model **v** = (1), lim_*n*→∞_ *ζ*_**v**_ (*m*) is the Euler-Riemann zeta function (Abramowitz and Stegun, 1972). It is well-known that lim_*n*→∞_ *ζ*_(1)_(2) = *π*^2^/6, lim_*n*→∞_ *ζ*_(1)_(3) = 1.202057, and lim_*n*→∞_ *ζ*_(1)_(4) = *π*^4^/90. Therefore

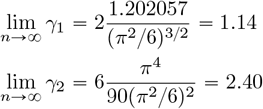

These results suggest that, asymptotically, *L* does not follow a normal distribution, since a normal distribution has both skewness and excess kurtosis equal to zero. This is somewhat surprising, as the Central Limit Theorem (Feller, 1968) commonly applies to the sum of a large number of independent variables with finite means and variances—such as *K*, which is proportional to *L* (in expectation) and is the sum of negative binomial variables (Griffiths, 1982; Durrett, 2008; Fu, 2025a). The failure of skewness and kurtosis to diminish to zero arises from the convergence of *ζ* _(1)_ 2 with sample size. In other words, the asymptotic finite variance of *L* prevents it from approaching normality.

Fig. 4 plots the dynamics of skewness and kurtosis of *L* for both the population of constant effective size and the fast growing population. Under the model **v** = (1), it can be seen that both skewness and kurtosis decline rapidly with *n* and when *n* = 20 (3 in log scale), *γ*_1_ = 1.190 and *γ*_2_ = 2.550, which are already rather close to their asymptotic values1.14 and 2.40 respectively. While under the model 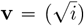 visible decline of both *γ*_1_ and *γ*_2_ towards zero with increasing sample size are observed.

**Fig. 4.**
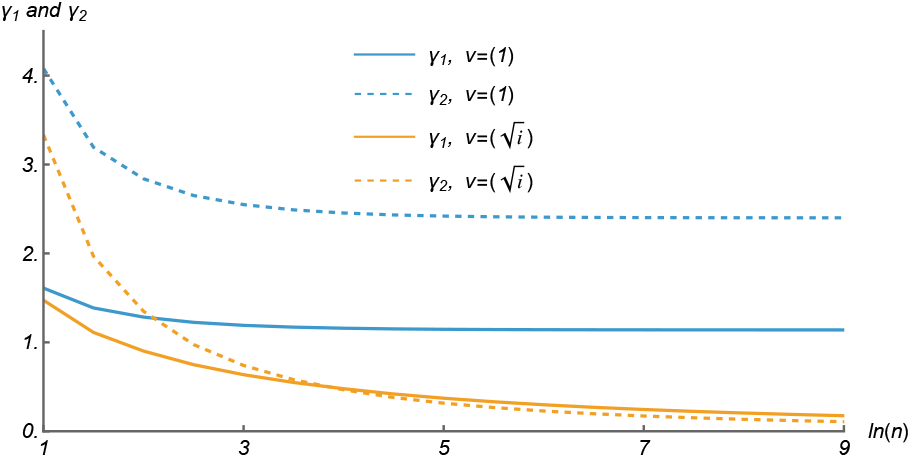
The Skewness and Kurtosis of *L* under **v** = (1) and 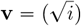 for sample size ln(*n*) ⩽ 9.

For a growing population with **v**, *ζ*_**v**_(2) increases faster with sample size than either *ζ*_**v**_(3) or *ζ*_**v**_(4), thus both *γ*_1_ and *γ*_2_ are generally smaller than those under the model **v** = (1). If there is an integer *c* such that for all *i* > *c*

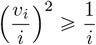

or equivalently 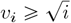, then

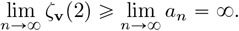

Therefore under the model 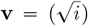 both the skewness and kurtosis approach those of normality with the increasing *n*. Since equality of finite moments is only a necessary condition, whether that *v*_*i*_ increases at speed of 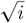 is suf-ficient for *L* to approach normality remains to be proven formally.

To inspect visually the difference between the density of *L* and that of a normal distribution, we used coalescent simulations to examine the density of the standardized *L*, which is

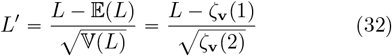

Fig. 5 shows that under the model *bv* (1), the densities for sample size 50 and 1000 differ little, both are far from that of *N* (0, 1). For the fast growth population model 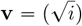, densities are much closer to the normality and there is a visible improvement with large sample size. For the rapid expansion model 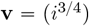 the agreement with *N* (0, 1) is further improved for both *n* = 50 and *n* = 1000 . These results also confirm the patterns in Fig.4 where the skewness and kurtosis differ little for ln(50) = 3.9 and ln(1000) = 6.9 under the model **v** = (1), but much smaller and yet appreciable difference between the two sample sizes under the model 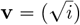.

**Fig. 5.**
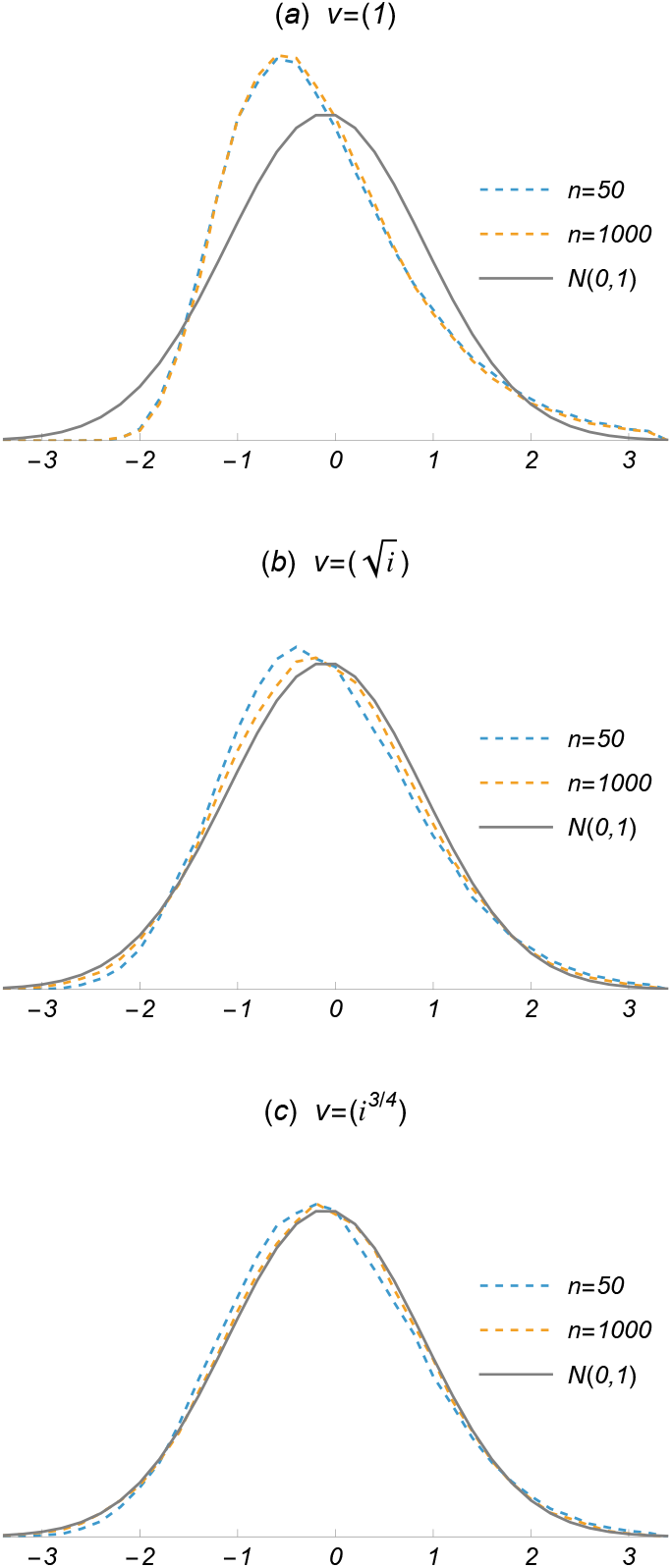
Estimated densities of *L*^′^ ( defined by Eq.(32) ) for two different samples sizes and under three constant-in-models (a) **v** = (1), (b) 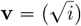 and (c) 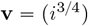. Each estimated density is from coalescent simulations with 10^5^ replicates.

**Fig. 6.**
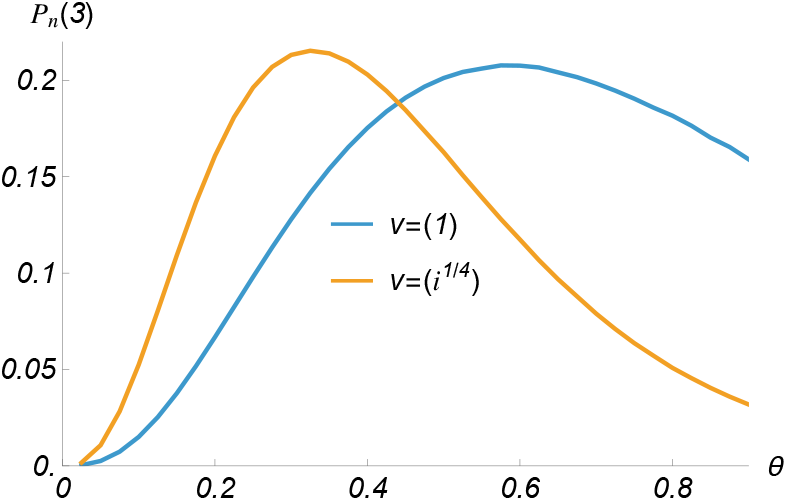
The likelihood of *θ* for *K* = 3 and *n* = 100 under two constant-in-state models.

## 5. Moments of *K*

The number *K* of mutations in the sample genealogy is ubiquitous. Its first and second moments were obtained by Watterson (1975), but its skewness and kurtosis were only recently explored by Fu (2025a) using its moment generating function. However, this standard approach is laborious and difficult to generalize to higher moments.

Alternatively, the power moments of *K* can be obtained using the argument of conditional expectation. Conditioning on the value of *L, K* is a Poisson random variable with parameter *Lθ*, where *θ* = 4*Nμ*, with *N* being the effective size of the reference population and *μ* the mutation rate per sequence per generation. Therefore, according to Haight (1967),

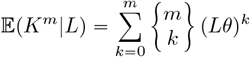

where 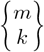 is Stirling numbers of the second kind (Abramowitz and Stegun, 1972). Thus the unconditional power moment of *L* is

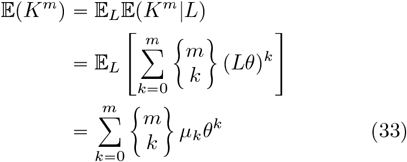

which can be computed numerically for any *m* since *μ*_*m*_’s are now available. Utilizing the concise expressions of *μ*_*m*_ in Eq.(19), it is straightforward to show that

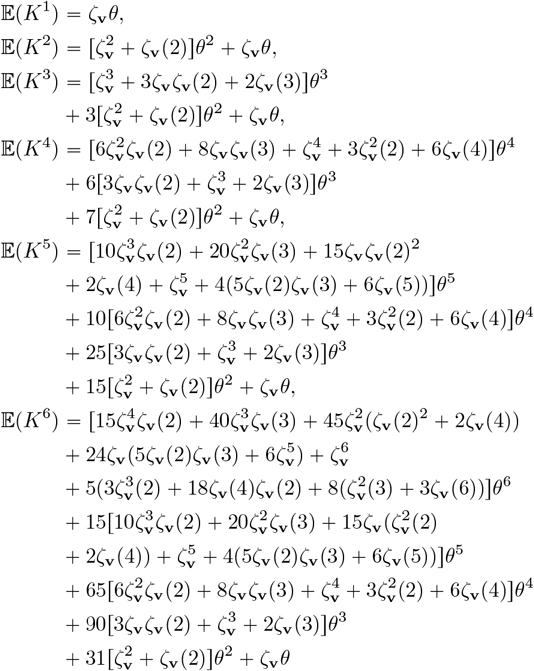

where 𝔼 (*K*^5^) and 𝔼 (*K*^6^) are unknown before while the 𝔼 (*K*^4^) in Fu (2025a) ( Eq.(38)) had some typos. Similar to Eq.(20), the central moments of *K* can be obtained as

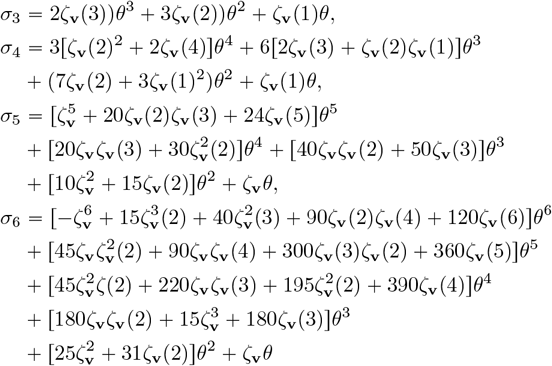

where *σ*_3_ and *σ*_4_ have been derived by Fu (2025a).

## 6. Distribution of *K*

The power moments of *L* can also be utilized for computing the probability of *K*. This is an area of little progress for decades until recently Fu (2025b) under the constant-in-state model. For the purpose of evaluating the probabilities of different *K*’s under a given *θ*, the sampling formula by Fu (2025b) is preferred, but the approach we present here has some advantages to warrant its existence.

Since *K* conditional on *L* is a Poisson variable with parameter *Lθ*, therefore

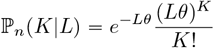

The unconditional probability of *K* mutations is thus

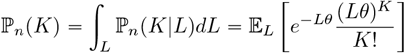

With Taylor expansion of *e*^−*Tθ*^, it leads to

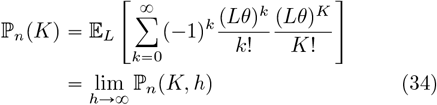

where

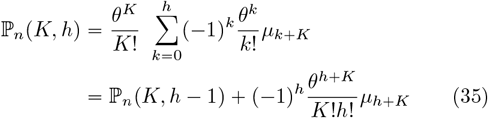

It is only necessary to evaluate ℙ*n*(*K, h*) for a sufficiently large *h* to satisfy a given accuracy, which can be specified with the relative error

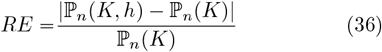

where ℙ_*n*_ (*K*) is the true probability of *K*, which can be computed using accurate method such as Eq.(11) of Fu (2025b).

Table 1 shows the minimum *h* for achieving 1% or smaller error for evaluating ℙ_*n*_(*K, h*) for some cases of *K*’s for two different values of *θ* under the model **v** = (1) and 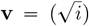. It is clear that the new approach is capable of obtaining accurate estimate of ℙ_*n*_ (*K*) efficiently as long as *K* is not large. In general for a given accuracy, the required *h* increases with both *θ* and *K*. It is also found that using relative change as the measure of accuracy is also effective, which is defined by replacing ℙ_*n*_(*K*) in RE by ℙ_*n*_(*K, h* − 1). That is

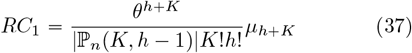

One advantage of the new method is that *μ*_*m*_’s only need to be computed once for a given sample size, so the probability under different *θ* can be evaluated easily. This suggests the new method has some advantages when the likelihood of *θ* is the focus. Fig.6 gives examples of like-lihood evaluation for *K* = 3 under two constant-in-state models.

However, the new method becomes inefficient when *K* is large and with increasing *θ* despite that ℙ_*n*_ (*K, h*) is guaranteed in theory to converge to ℙ_*n*_ (*K*) . Also since *μ*_*m*_ increases exponentially when *m* is large, which is prone to rounding errors and may eventually exceeds upper bound of number in a computational environment, the method is thus not recommended for general use for *θ* ≫ 0.7.

## 7. Probability of a single mutation of certain size

The approach presented in the previous section for computing ℙ_*n*_ (*K*) can be adopted in principle for computing the probabilities of patterns of mutations. By a pattern of mutations, we mean the specification of not only the number of mutations, but also the size of each mutation, which is conveniently represented by a multiset

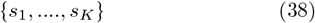

where *s*_*k*_ is the size of the kth mutation (order of mutations is immaterial and is for convenience only). This is an important area that deserves much attention. To start, we consider the the probability of a single mutation of certain size. That is,

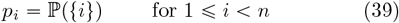

This probability is of interest since it is the dominant pattern of mutations in a sample from a relatively small locus.

Since a mutation of size *i* can only occur on one of the branches of size *i*, which total length is *L*_*i*_, the probability of the mutation pattern {*i*} conditional on coalescent times (*t*_*k*_, *k* = 2…, *n*) is

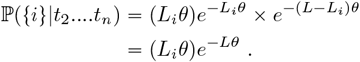

That is, it is the joint probability of the event that there is one mutation of size *i* and the event that there is no mutation on any branch of size *j* ≠ *i*. Consequently the unconditional probability of {*i*} is

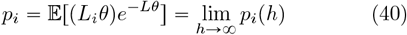

where

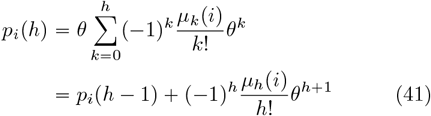

Due to Eq.(28), we have as expected

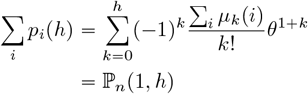

Similar to the case of ℙ_*n*_ (*K*), one can satisfy a given accuracy by evaluating *p*_*i*_ (*h*) for a sufficiently large *h*, and the accuracy can be measured by the relative change:

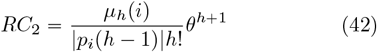

Fig. 7 plots *p*_*i*_ (*h*) against *h* for several values of *i* under several scenarios for a sample of size 100. It is clear that *θ* is the dominant factor in determining the pattern of *p*_*i*_ (*h*) . With smaller *θ, p*_*i*_ (*h*) becomes stable around *h* = 7, while for larger *θ* = 0.5 there is still visible variation around *h* = 13. These patterns follows similar ones for ℙ_*n*_ (*K, h*) but in general smaller *h* is required to evaluate *p*_*i*_ (*h* ).

**Fig. 7.**
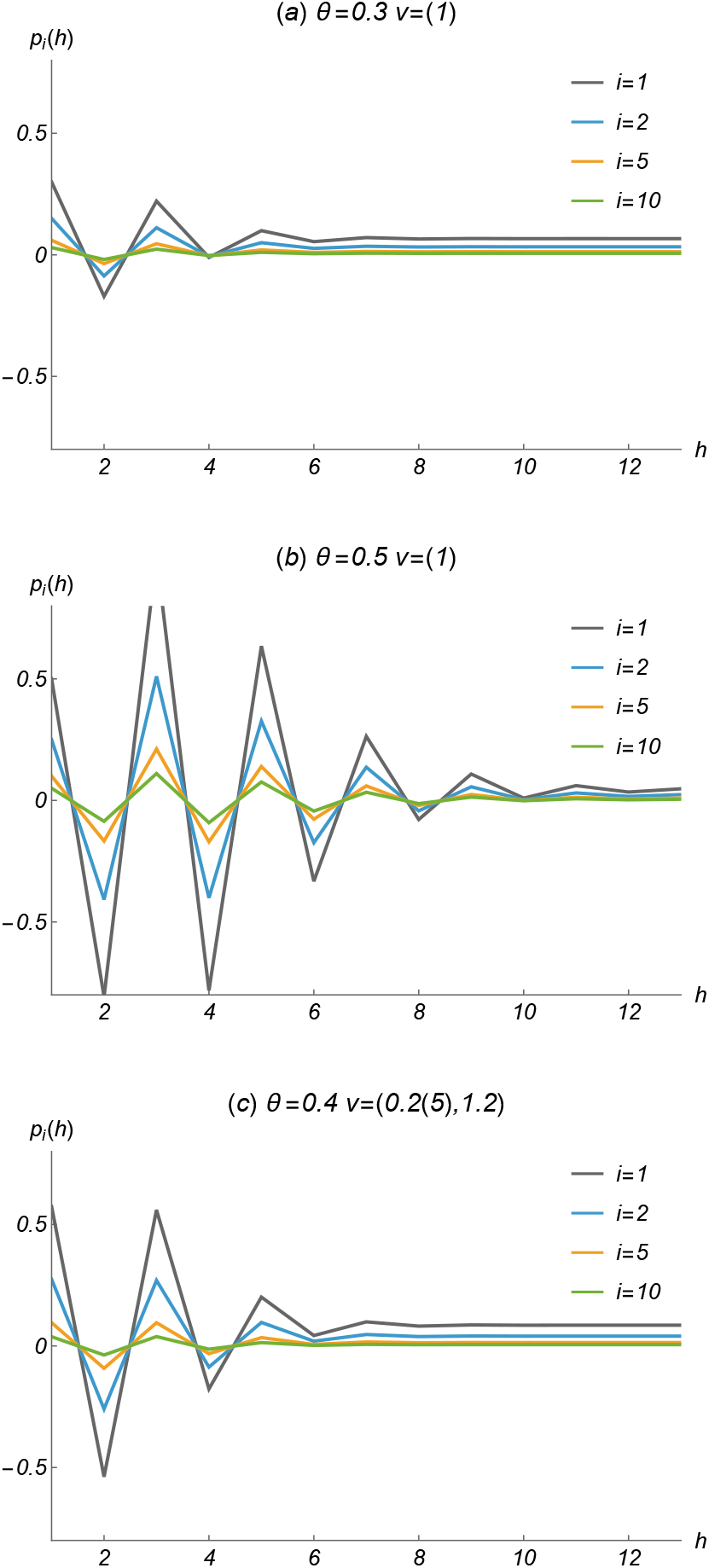
Effects on the estimate of *p*_*i*_ by the number of terms in Taylor expansion (41) for constant population (a) and (b), and a constant-in-state model (c). Sample size n=100 for all three scenarios.

Fig. 7 also shows an example of *p*_*i*_ *h* under an expanding population (case c). The pattern appears to be consistent with the expectation that it should be somehow between those of case a and b. It appears regardless of model specification **v** that *θ* is the primary factor determining the *h* for achieving a given accuracy. For *θ* ⩽ 0.5, it appears that *h* = 15 is generally adequate. Similar to the recommendation made in the previous section for computing ℙ_*n*_(*K*), it is not recommended to use the method for *θ* ≫ 0.7.

## 8. Discussion

This study systematically explores the power moments *μ*_*m*_ and *μ*_*m*_ (*i*), leading to a deeper understanding of the properties of *L* and its applications under the constant-instate model. That *L* does not approach normality under the constant-size Wright-Fisher model is unexpected, as asymptotic normality has been established for *K* and the number of alleles in the sample. The root cause of this non-normality is the finite variance of *L* as the sample size increases, which can be overcome if the population grows sufficiently fast.

The closed-form formula for *μ*_*m*_ is related to the integer partitions of *m* and the number of permutations of cycle types associated with each partition. This provides another example of the importance of cycle-type permutations in population genetics, which appear in Ewens’ sampling formula (Ewens, 1972; Karlin and McGregor, 1972) and more recently in the sampling formula for *K* (Fu, 2025b).

The power moments *μ*_*m*_ also lead to a novel method for computing the power moments of *K*, which have so far been limited to the first few using the standard method based on its moment generating function. These power moments are also useful for computing the probability of *K*, providing an alternative to existing methods with the added advantage of efficiently evaluating multiple values of *θ*.

A particularly promising application of *μ*_*m*_ and *μ*_*m*_ (*i*) is in computing the probabilities of single mutations of certain sizes. Such probabilities are essential for inferring population dynamics. Even more importantly, this approach has the potential to be extended to compute the probabilities of patterns involving two or more mutations.

One caveat of using *μ*_*m*_ and *μ*_*m*_ (*i*) is that they increase rapidly with *m*, which makes them prone to rounding errors for large *m*. Therefore, caution should be exercised when applications require very large *m*. For human genomes, with the exception of recombination hotspots, the assumption of no recombination is likely valid for chromosomal segments up to approximately 500 bps, and may be reasonable for segments up to 1000 bps. These correspond to *θ* ⩽ 0.35 and *θ* ⩽ 0.70, respectively, based on our analyses of the 1000 Genomes Project data, which are well within the applicable range of *θ* for our novel methods for computing the probability of *K* and the probabilities of mutation patterns.

### Software

A Java package for computing the moments of *L*, the probability of *K* and the probability of a single mutation of certain size is available upon request.

